# Spacing spheroids, curl-up and coiled neuritic plexus of the sacral spinal cord of aged dog: Hypothesis based on continued NADPH diaphorase histological observations

**DOI:** 10.1101/2020.03.31.018234

**Authors:** Yunge Jia, Yinhua Li, Wei Hou, Huibing Tan

**Author notes:** **Correspondence:** Department of Anatomy, Jinzhou Medical University, Jinzhou, Liaoning 121001, China. The first two authors make equal contributions to this work.

## Abstract

NADPH diaphorase (N-d) neurons distribute in spinal cord and function for visceral sensation and autonomic regulation. N-d positive neurons innervate pelvic organs. In previous investigation, we report that aging-related N-d body (ANB) in the lumbosacral spinal cord in aged rat and megaloneurite in the sacral spinal cord in aged dog. This article was a continued data report of aging-related N-d alterations in aged dog. N-d positivity in aged spinal cord has revealed a certain of morphological profiles in the spinal cord of several species. However, we still found some denoted N-d neurodegenerative changes that we failed to notice in our previous studies when re-examination of the sacral spinal cord of aged dog. In the horizontal section, spacing spheroids in the superficial laminae of the dorsal horn, curl-up and coiled neurites in the intermediate zone were detected in the sacral spinal cord. The ANB and vacuolar neurite were also detected. Vacuolar degeneration also occurred in the dorsal ganglia at the sacral segment. The curl-up and coiled neurites did not occur in the lumbothoracic segment, but the ANB and vacuolar neurite were scatteringly detected in in the lumbothoracic segment of aged dog. The results suggested that the N-d sensory inputs interrupted and disconnected with integration of autonomic centers and output circuits of regulating urogenital organs during the aging. These specialized profiles were speculated that the N-d neurite deterioration of visceral sensory circuit implicated dysfunction of pelvic organs in the aging. Megaloneurite and fiber dilation may make backward reasoning to N-d fiber architecture under normal condition.

## INTRODUCTION

Urinary incontinence is clinical epidemiological health issues of older adults[1] Constipation is also common in adults older than 60 years [2]. Generally, constipation is considered for the peripheral enteric neurodegeneration[3, 4]. Erectile dysfunction in men is also a sign of an underlying aging condition[5]. The sensation of the aging spinal cord could be changed in the singling circuitry in the dorsal horn of the lumbosacral segment [6]. The impact of aging on the cellular composition of the spinal cord reveals aging-associated changes in the numbers of neurons and non-neuronal cells[7].

The sacral spinal cord innervates of the urogenital organs [8, 9]. The relevant cell bodies of autonomic neurons distribute in the dorsal horn, intermediolateral nucleus (IML), dorsal commissural nucleus (DCN). In the dog [9, 10] and monkey[11], autonomic neurons also locate in the anterior horn. Many neuropeptides distribute in these neurons[12, 13]. The NADPH diaphorase (N-d) is also important neuronal enzyme identical to nitric oxide synthase (NOS) distributed in the autonomic spinal cord[14–16], although some experiments may not completely reach the consistent conclusion [17–20]. Up to date, the N-d is also commonly used to investigate the neurodegenerative diseases[21–23]. Aging-related N-d alterations have been studied in the sacral spinal cord in the rat[18], dog[24, 25], pigeon[26] and monkey[27]. In these experiments, the aging-related N-d alterations show the diversity of the neuropathological profiles. Recently, by using N-d histology, we re-examined the sacral spinal cord of aged dog. We continued to upgrade some previous neuropathological study in this brief communication.

## MATERIALS AND METHODS

### The tissue preparation

The spinal cords of aged dogs (Canis lupus familiaris, less than 2-year-old, n=5; more than 8-year-old, n=6) of both sexes was legally obtained from Department of Surgery Experimental Animal Facility of Jinzhou Medical University (Jinzhou, China). The experimental protocol was approved by the Ethics Committee on the Use of Animals at Jinzhou Medical University.

The dogs were deeply anesthetized by intravenous injection of sodium pentobarbital (overdosage of 60 mg/kg body weight). The protocol of perfusion was slightly modified according to the previous experiments[28, 29]. Briefly, the dogs were trans-cardially perfused through the aorta with normal saline, followed by 4% paraformaldehyde in 0.1 M phosphate buffer (PB; pH 7.4). The tissues of the spinal cord were rapidly obtained and immersed in the same fixative as in the perfusion for at least 6 hrs. at 4°C, and transferred to 30% sucrose in 0.1 M PB (pH 7.4). Frozen sections were cut transversely or horizontally at 40 μm on a cryostat (Leica, German).

### NADPH diaphorase histochemistry

N-d enzyme-histology was performed in free-floating method. In this procedure, sections were incubated for 5 min in 100 mM sodium phosphate buffer (PBS, pH 7.4) followed by incubation in phosphate buffer (PB, pH 7.4) with 1 mM beta-NADPH (Sigma, USA), 0.5 mM nitrotetrazolium blue (NBT, Sigma, USA) and 0.3% Triton X-100 at 37℃ for up to 2-4h. Sections were rinsed with PB, distilled water, dehydrated in a graded ethanol series, and were coverslipped with mounting medium.

### Quantitative microscopy and Statistical analysis

Sections were observed under the light microscope (Olympus BX53 microscope, Japan). Images were captured with a DP80 camera. Sections from all spinal cord levels in each animal were quantitated using Olympus image analysis software (Cellsens Standard, Olympus). Mean value was presented as mean ± SEM. Statistical significance was accepted with p≦0.05.

## RESULTS

We first examined the coronal transection of the spinal cord in the young adult and aged dog. N-d positive neurons and fibers distribute in the dorsal horn and around the central canal, the intermediolateral column (IML), the Lissauer’s tract (LT) and lateral collateral pathway [30] in sacral segments in young adult dog (Figure 1). This was consistent with the previous report [24, 31]. A few of N-d fibers projected ventrolaterally to the IML region (Figure 1 B). The neurons also distributed in the ventral horn [24], which is a different feature from rats. The number of cell bodies in the ventral horn was relative much numerous than that in the dorsal part of the spinal cord, but the fibers and their varicose nature in the ventral were not quite evident compared with the dorsal part of the spinal cord. In previous studies, we found that megaloneurite of N-d positivity is the major aging-related alteration in the sacral spinal cord of aged dog[24]. The megaloneurite distributed in the LT, LCP and DCN in aged dog, compared with the thin fiber bundles of axons in the LT, LCP and DCN of young adult dog. Our present finding of the megaloneurite was consistent with the previous study. In coronal sections of the sacral spinal segments, a cluster of N-d stained spheroids were noted in the intermediate lateral region (Figure 2). Among the spheroids (open arrow), segment of thick neurite (curved open arrow) was detected in Figure 2D. In following horizontal sections in Figure 6, the cluster of spheroids were corresponding to the location of the coiled neurites. In order to further demonstrate the orientational fiber-architecture at the sacral level, an orientation drawing of horizontal sections in the sacral segment (Figure 3). Next, we showed at least three levels from which following figures would demonstrate the spacing mini spheroids and coiled neurites.

**Figure 1.**
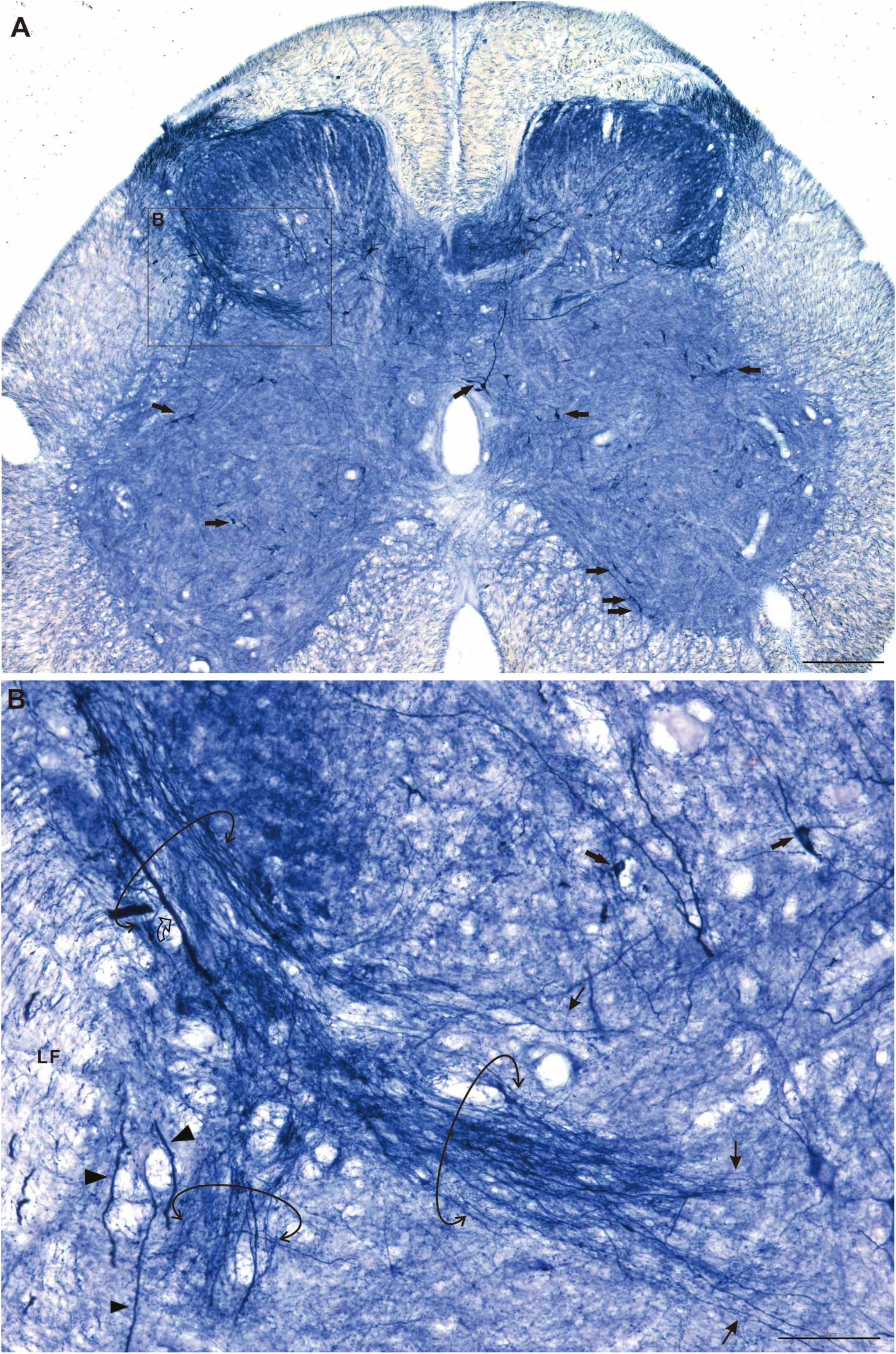
N-d staining in the sacral spinal cord of young adult dog. Arrow: neuron. Circle arrow: fiber tract; Arrowhead: thick fiber; thin arrow: thin fiber; curved open arrow: thick fiber. Bar A= 200μm and B = 50μm.

**Figure 2.**
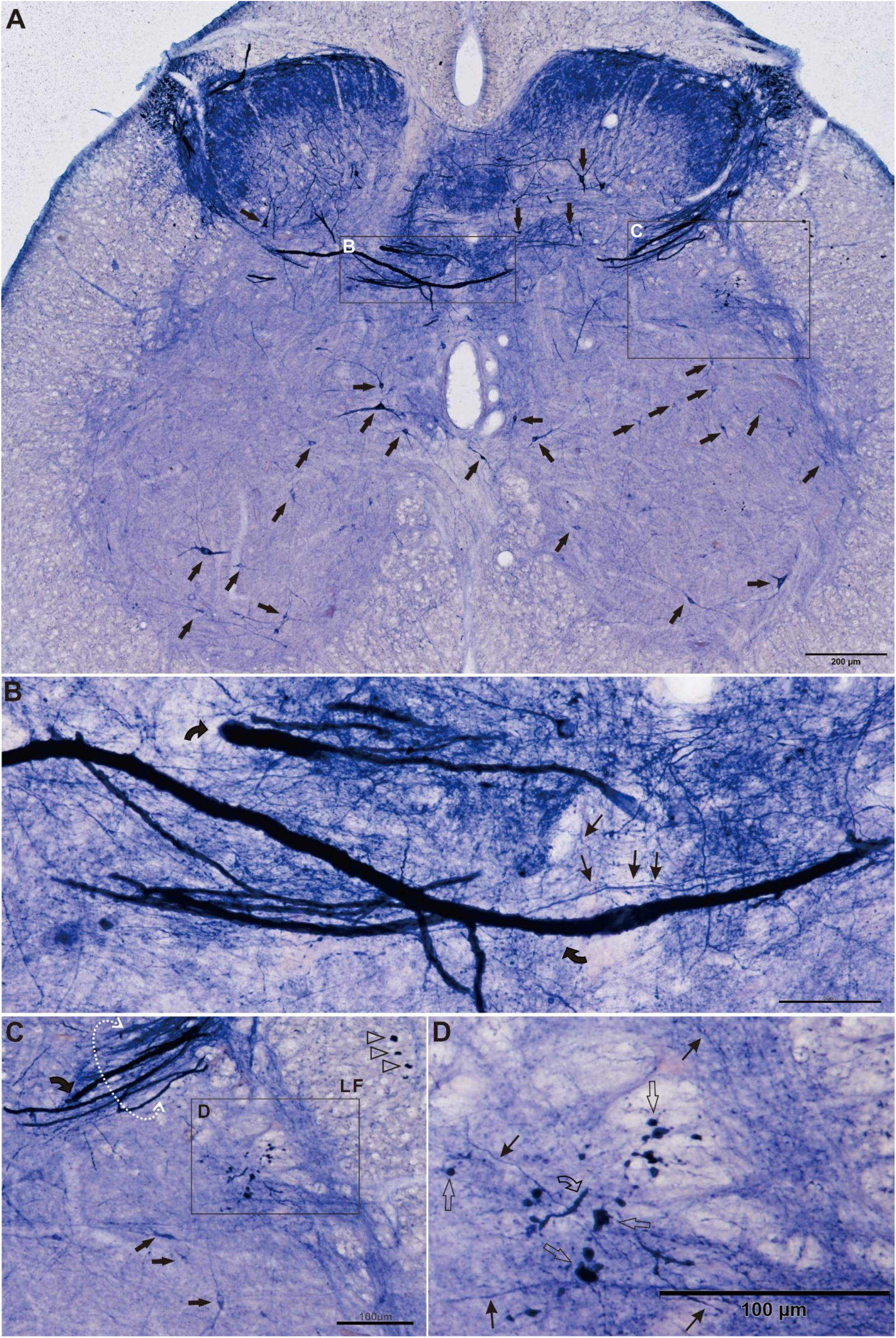
Megaloneurites (A and B) and spacing spheroids (C and D) in the sacral spinal cord of aged dog. Arrow indicated neuron. B showed megaloneurites (curved arrow) and thin fiber (thin arrow) from A. Transverse of megaloneurites (open arrowhead) in the lateral funiculus (LF). Circle arrow indicated lateral collateral pathway(C). Spheroids (open arrow) and segment of thick neurite (curved open arrow) detected in D. Bar in A=200μm, B=50μm, C and D =100μm.

**Figure 3.**
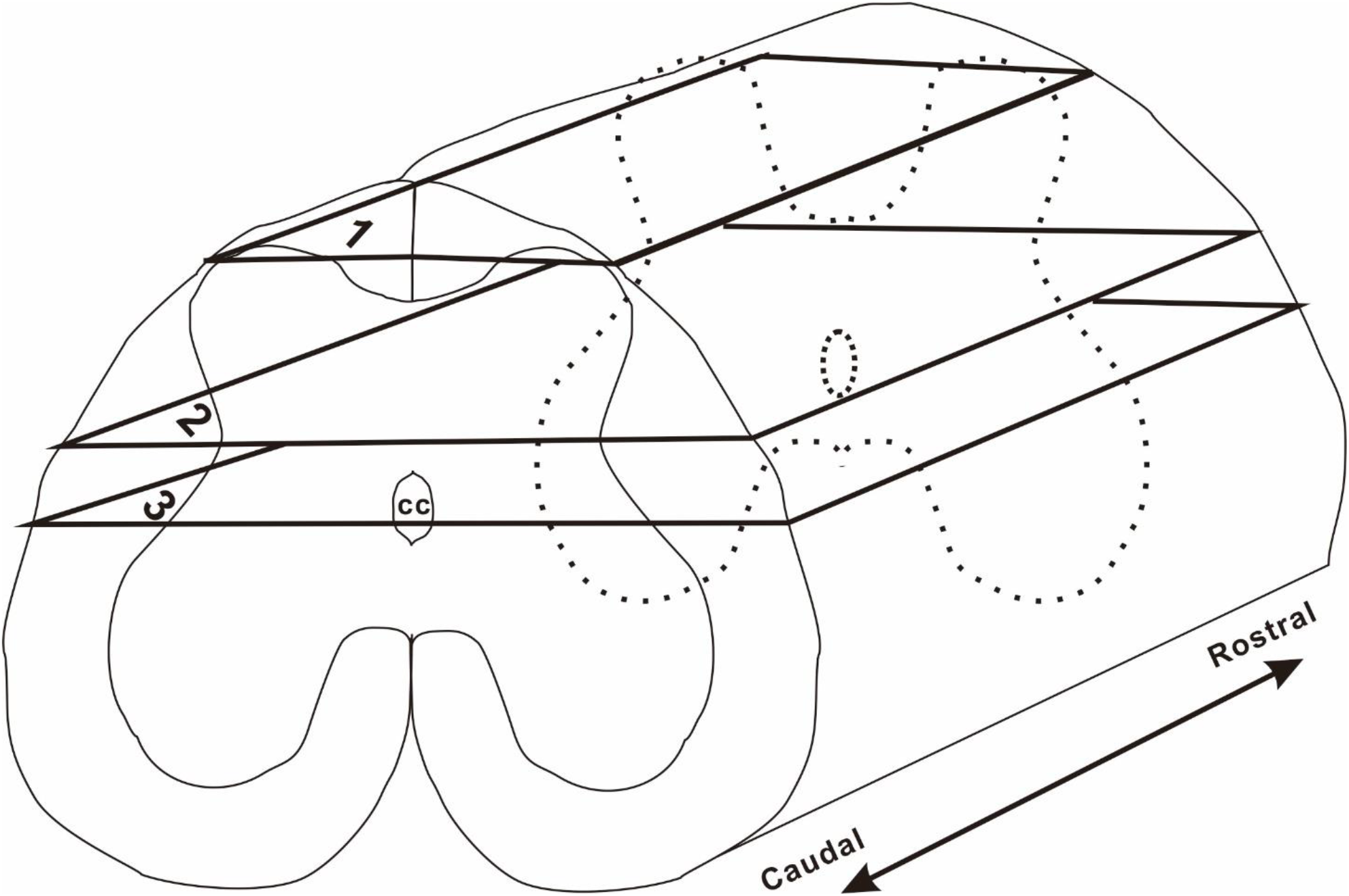
Orientation drawing of horizontal sections in the sacral segment showing three levels from which following figures would demonstrate the spacing mini spheroids and coiled neurites. cc indicated central canal; 1 represented the level for the dorsal marginal laminae, 2 represented the level for the dorsal commissural gray; 3 for the level cross the central canal.

As indicated drawing level 1 in Figure 3, Figure 4 showed neuropil mini spheroids distributed in the dorsal marginal laminae in the dorsal horn of horizontal section in aged dog. Definitely, high power magnification of the Figure 4 B revealed the megaloneurites (circle double arrow) vs. thin fibers (arrowhead). The maximum of the diameter of megaloneurites reached 26.01μm in the histogram. The average diameter of the megaloneurites was significantly thicker than that of regular thin fibers(p<0.05). Some thick fibers occurred in LT (circle arrow in Figure 4C). The diameter of the fibers gradually increased the thickness of the neurites from the surface to deeper parenchyma along the LT. In the Rexed laminae I, some mini spheroids occurred in the neuropil of the marginal zone of the spinal cord. Some of the mini spheroids occurred along the thin fibers. There were also more relative larger size spheroids distributed in the same regions. Next, another three examples of the similar locations were presented to show large size spheroid and swelling (dilating) neurites in the dorsal marginal laminae.

**Figure 4.**
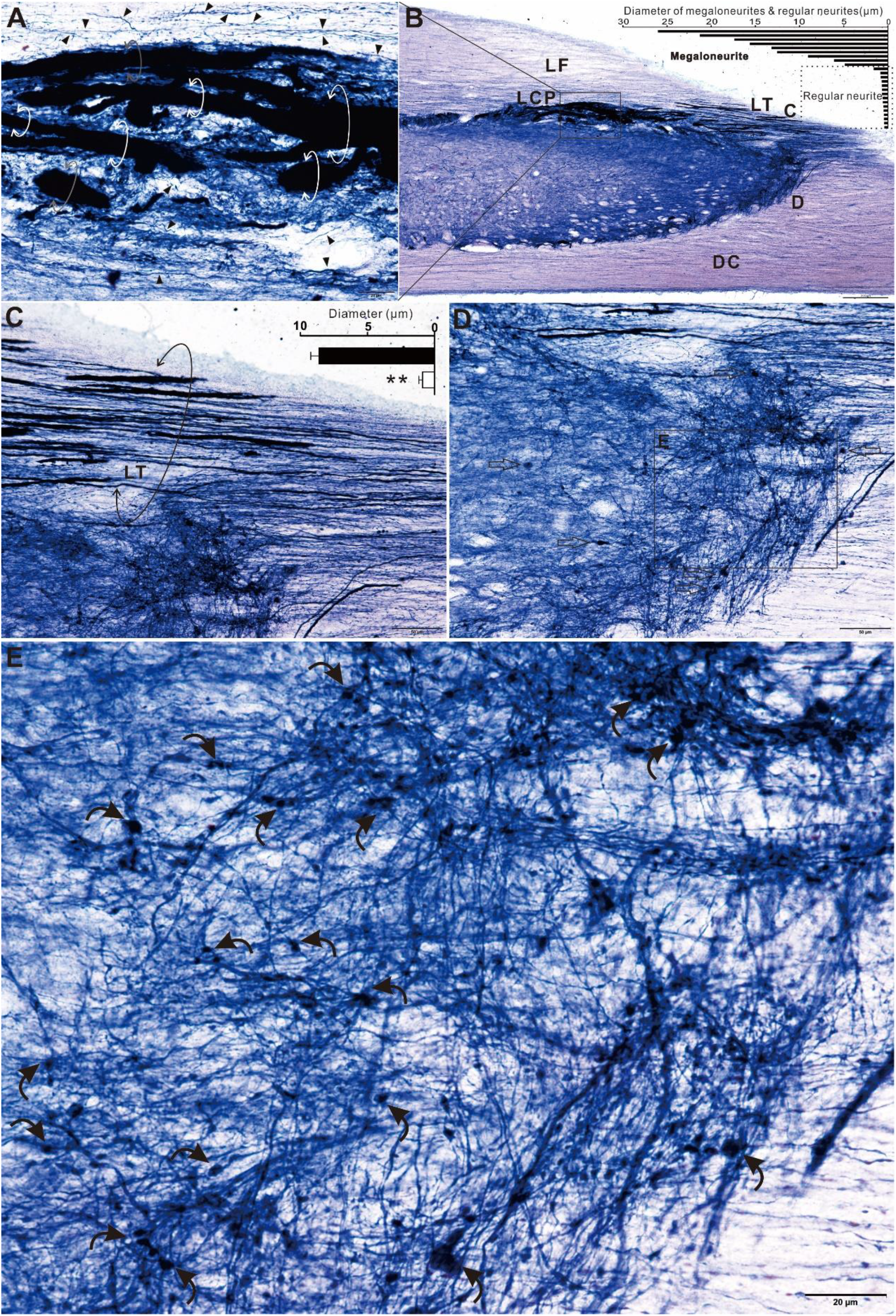
Neuropil mini spheroids distributed in the dorsal marginal laminae in the dorsal horn of horizontal section (aged dog, indicated drawing level 1 in Figure 3). A: High power magnification of the B indicated the megaloneurites (circle double arrow) vs. thin fibers (arrowhead). B: Low magnification for A. The histogram (right up corner) indicated diameter of megaloneurites and thin fibers measured from A. LF: lateral funiculus; LCP: lateral collateral pathway; LT: Lissauer’s tract; DC: dorsal column. C and D indicated the regions for Magnified image C and D. Some thick fibers occurred in LT(circle arrow in C). Open arrows in D indicated some spheroids. The localization of E in D was the marginal zone (Rexed laminae I) of the spinal cord. Some mini spheroids distributed in the neuropil. Bar in A and E =20μm, B=200μm, C and D =50μm.

We added one horizontal section Figure 5-1 similar location as Figure 4 to demonstrate N-d longitudial neurites and neurodeeneration in the marginal laminae in the sacral dorsal of aged dog. The longitudual neurites varied from thin and thick fibers as well as megaloneurites. The pathway of the all neurites revealed fasciculus proprius and or cross segment funiculus. Figure 5-2 followed the same section of Figure 5-1 to review three microscope fields (A-D, E and F). Figure 5-2A revealed the megaloneurites(curved arrow) in the LT (circle arrow), lateral collateral pathway [30] [30] and medial collateral pathway (MCP). C showed that spheroids (open arrow) located in focal regions of swelling neurite. Curved thin arrow indicated mini spheroid. D also showed mini spheroids typically along the neurite. E showed two large spheroids between LT and marginal laminea. F showed inset of compound spheroid with relative shallow staining halo surround high intensity core. In order to show more detailed distribution of the enlarged spheroids. We presented magnified photo image of Figure 5C (supplementary figure 1).

**Figure 5-1.**
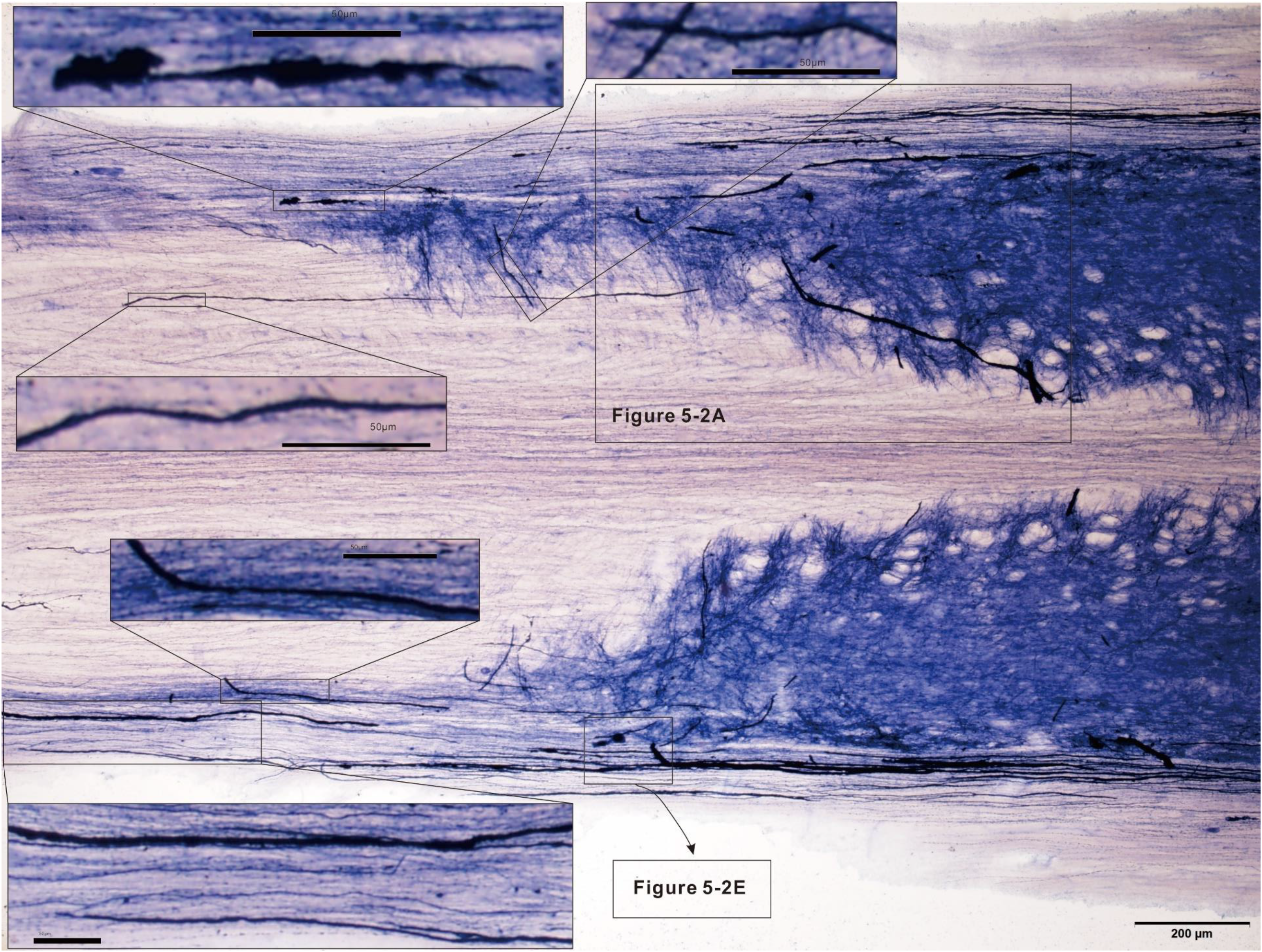
Horizontal section similar location as Figure 4 to demonstrate N-d longitudial neurites and neurodeeneration in the marginal laminae in the sacral dorsal of aged dog. Bar =200μm and inset =50μm.

**Figure 5-2.**
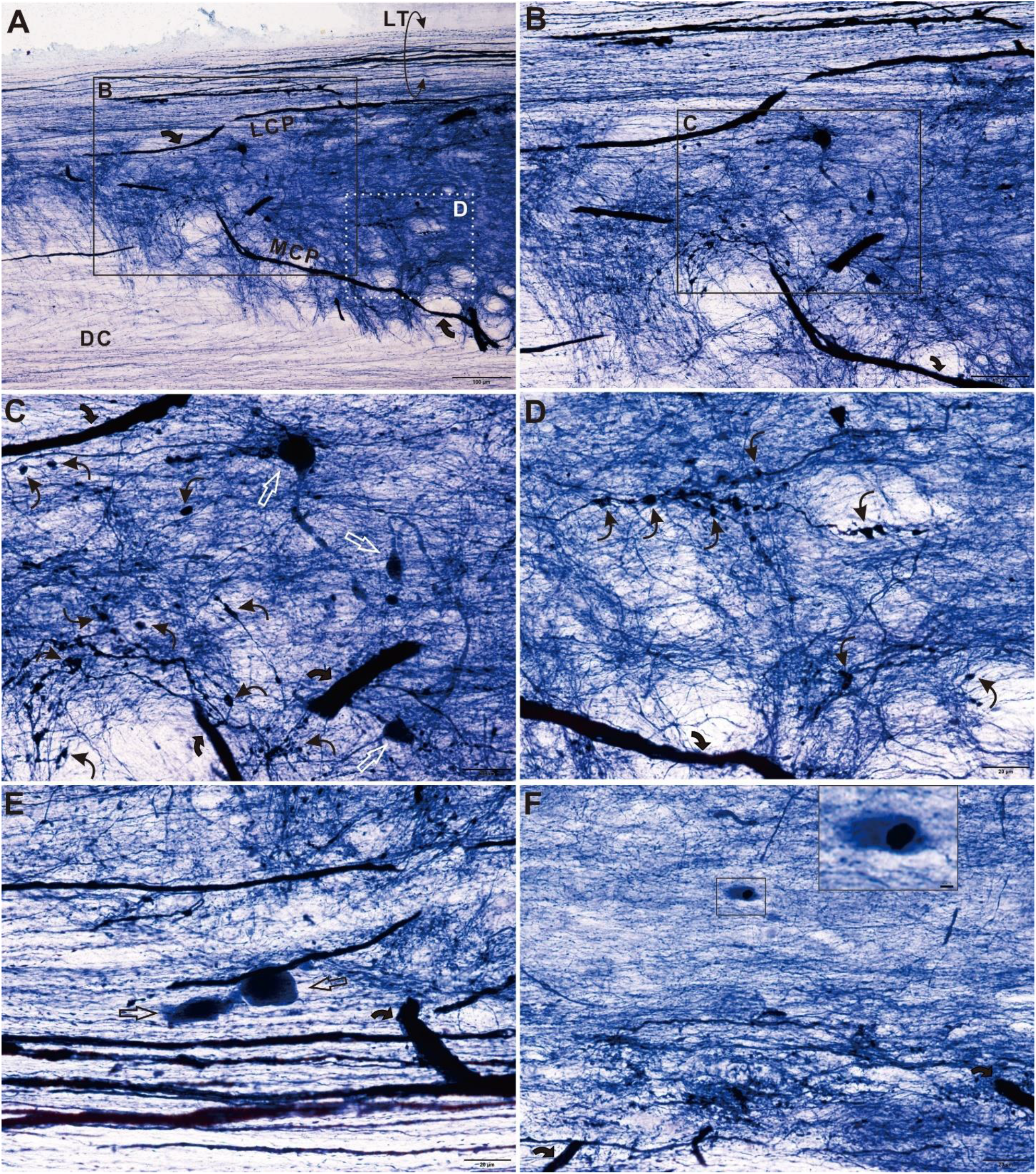
Continued demostration following Figure 5-1 with three microscope fields (A-D, E and F). A revealed the megaloneurites(curved arrow) in the Lissauer’s tract (LT, circle arrow), lateral collateral pathway [30] and medial collateral pathway (MCP). DC: dorsal colummn. B, C and D magnified from A. C showed that spheroids (open arrow) located in focal regions of swelling neurite.Curved thin arrow indicated mini spheroid. D also showed mini spheroids typically along the neurite. E showed two large spheroids between LT and marginal laminea. F showed inset of compound spheroid with relative shallow staining halo surround high intensity core. Bar A = 100μm, B=50μm, C-F=20μm and inset=2μm.

**Figure 6.**
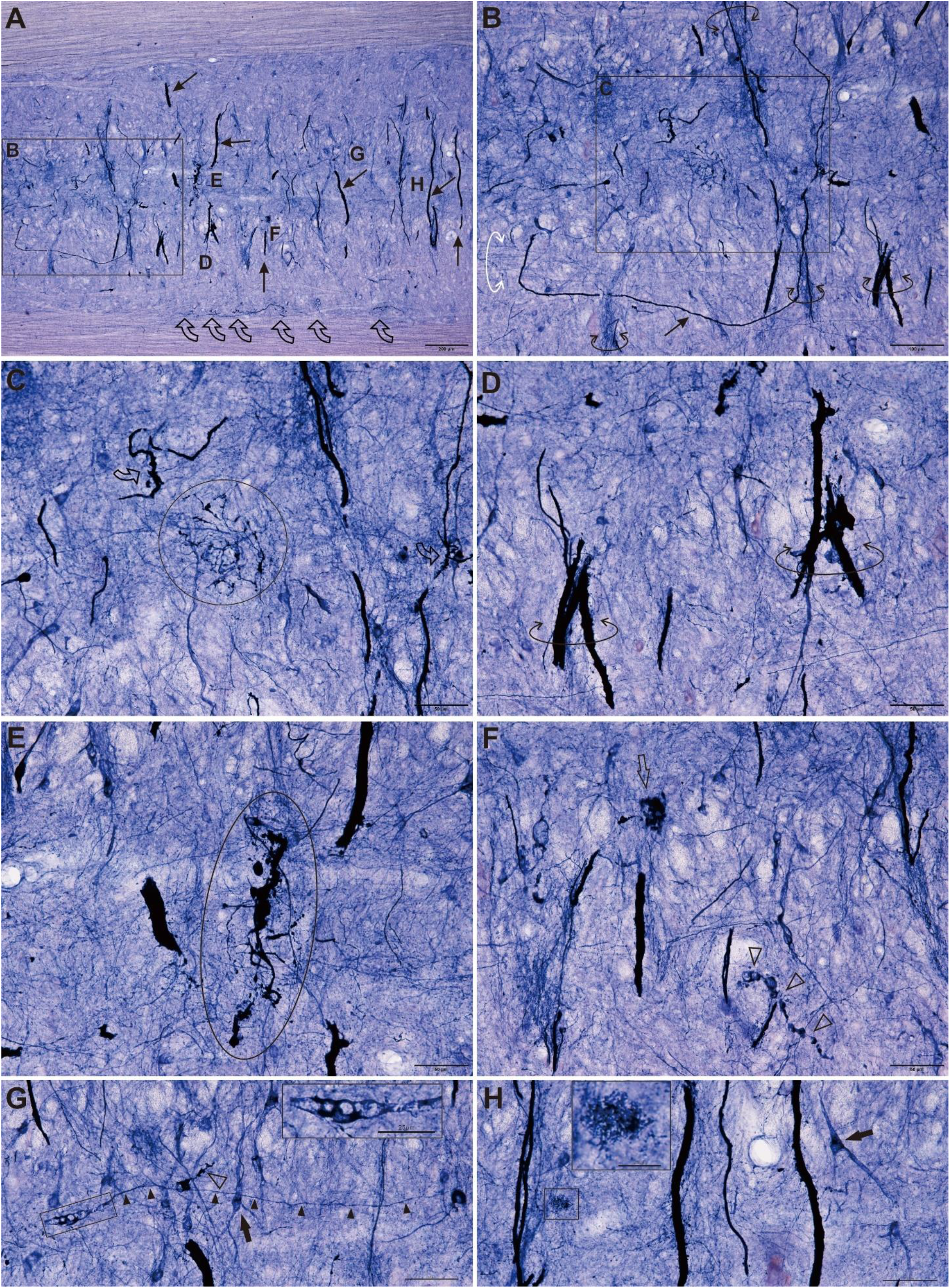
Horizontal section through the dorsal commissural gray indicated in drawing level 2 of Figure 3. A: Spacing arrangement of segments of megaloneurite (thin arrow) and curl-up neurite (curved open arrow). D, E, F, G and H labeled for following magnified images. B: Thin arrow indicated U-shape oriented neurite. White and black circle arrow indicated track megaloneurite and neurite in two oriented tracks. Coiled neurite (circle) and curl-up neurite (open curved arrow) in C. Circle arrow indicated segmental tracks of megaloneurites in D. E showed curl-up neurite (oval circle). Aging-related N-d body (open arrow) and vacuolar neurite (open arrowhead) in F. Inset in G showed vacuolar neurite with thin fiber(arrowhead). Arrow indicated neuron in G and H. Inset in H showed a patch of dots. Bar in A =200μm, B=100 μm, C-H=50μm and bar in inset =25μm.

According to Figure 3, Figure 6 was a section as the level 2 in the drawing illustration (Figure 3), which was a horizontal section through the DCN or term as dorsal commissural gray. Spacing arrangement of segments of megaloneurites revealed in the track of fiber bundles in an interval pattern. Some curl-up neurites revealed at bottom side of the Figure 5-2A. The curl-up neurites would be further demonstrated in another section indicated in Figure 6. The thick neurites and megaloneurites are a striking morphological feature for fiber architecture. We first report the findings in our previous investigation. Here, we chosen a U-shape oriented neurite in Figure 5-2B to demonstrate that neurol circuit based on the thick neurites and megaloneurite may also originate from an interval structure and may play important role in autonomic regulation. Acording to Figure 2 and Figure 3 to 5, the megaloneurites occurred in the LT, dorsal lateral funiculus, MCP and LCP. The aging changes of coiled neurite and curl-up neurite were detected in Figure 5C and E. The aging-related N-d body and vacuolar neurite were also detected in Figure 5-2F, G and H.

The major finding of the research was the spacing coiled neurites. Figure 7 showed a horizontal section in drawing level 3 of Figure 3 to show spacing coiled neurites along the intermediate lateral region in the sacral spinal cord of aged dog. The Figure 7-1A was the horizontal section through the central canal and the DCN. The spacing coiled neurites located in both sides of intermediate lateral regions (Figure 7-1 and 7-2). It was interesting that some coiled neurites apparently presented a circle or semi-circle format the diameter of which was quite consistent (Figure 7-1C and D). Semicircle arrow indicated medial to DCN fiber bundle, which showed circuit connection between the coiled neurites to DCN. Again, the vacuolar spheroid and regular spheroids were also detected in this section (Figure 7-2). In order to present detailed N-d positivity, we present two large format images which were the same photo image of Figure 6-1B and Figure 7-1C (Supplement 2 and Supplement 3). The coiled neurites did not be found in the thoracic segment of aged dog (data not showed).

**Figure 7-1.**
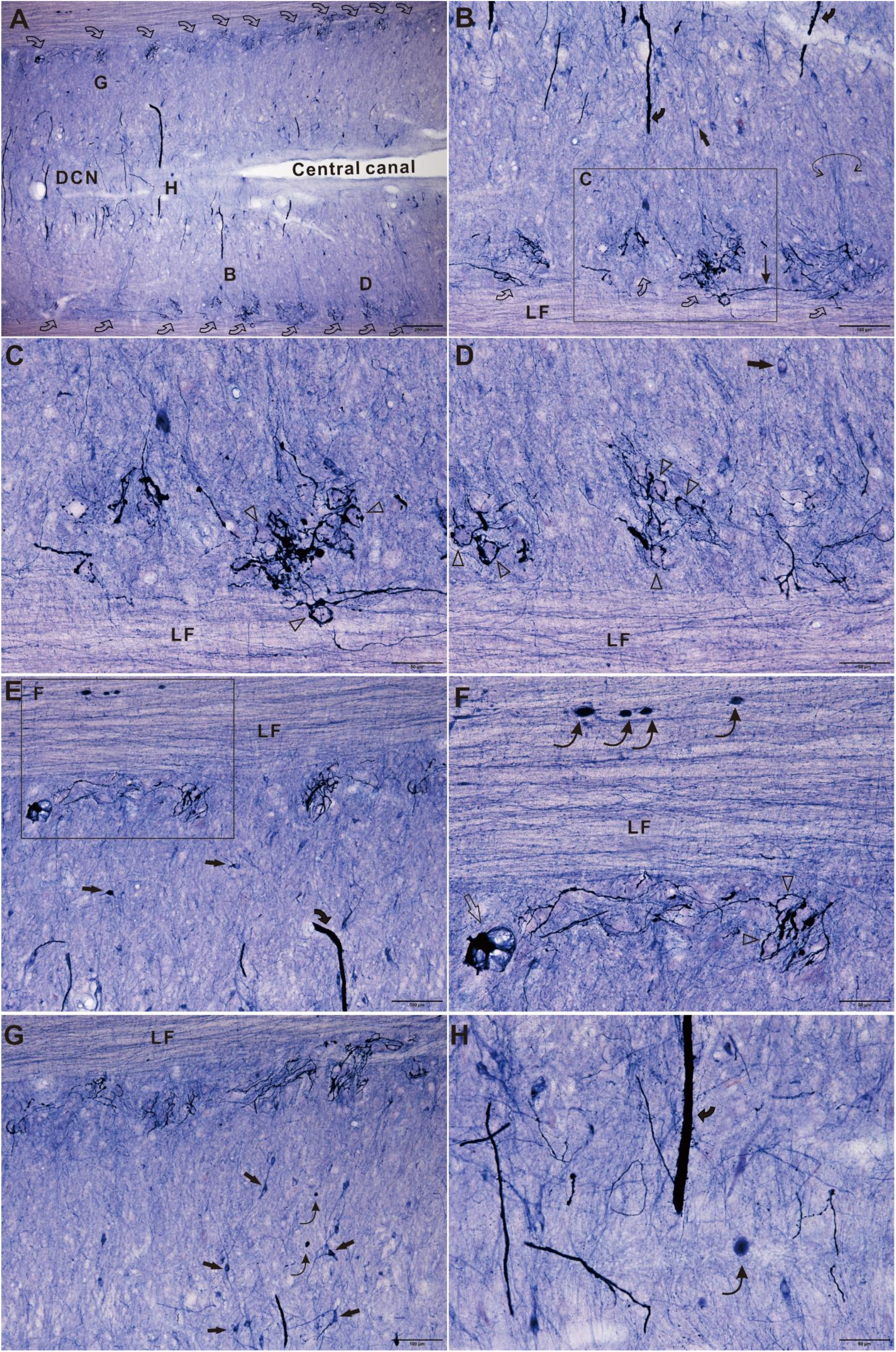
The section indicated in drawing level 3 of Figure 3 to show spacing coiled neurites along the intermediate region in the sacral spinal cord of aged dog. A: The horizontal section through the central canal and the dorsal commissural nucleus (DCN). Open curve arrow indicated spacing coiled neurites. B, D, G and H labeled for following magnified regions. B showed arrangement of coiled neurites (open curve arrow) and megaloneurite (curved arrow) as well as neuron (arrow) inside lateral funiculus (LF). Semicircle arrow indicated medial to DCN fiber bundle. C magnified from B. Some coiled neurites for circle or semi-circle the diameter of which was quite consistent (C and D). Bar in A =200μm, B=100μm, C and D =50μm.

**Figure 7-2.**
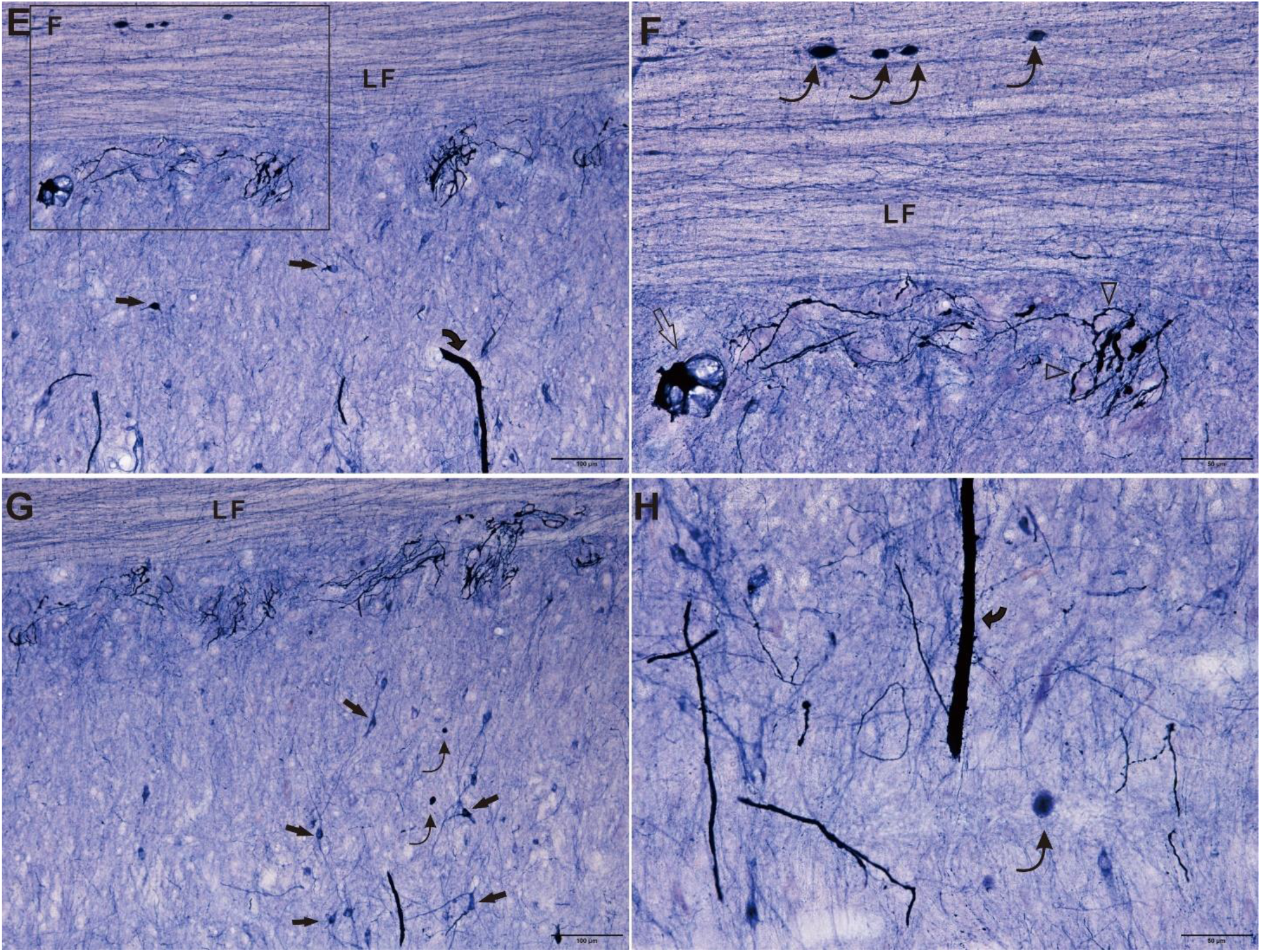
All photo images continued demonstration of Figure 6-1. E showed the other side of section from Figure 6-1A. F magnified from E showed there were spheroids (curved thin arrow) in the LF. Open arrow indicated a large vacuolar spheroid. G magnified image cross the central canal. Arrow indicated neurons. Curved thin arrow indicated spheroid. H showed region in the DCN. Bar in E=100μm, F-H= 50μm.

In our previous study, we report that the dystrophic neurite and megaloneurite also occur in the central canal[24]. Now, we reexamined the aging alteration. Figure 8 showed various diverse neurites in the horizontal section through the central canal. We demonstrated that some neuronal dystrophy around the central canal formed unique morphological profile. It was aberrant profile of dystrophic neurites, not uniform swelling, not identical megaloneurite and not simple spheroid as we report before. The Figure 8 B, there were at least three longitudinal oriented neurites revealed nearby ependymal cells, besides curl-up neurites. It was labeled with open arrowhead (dash line like fiber), arrowhead (varicose fiber) and finger point (thin fiber). Contrastingly, the curl-up neurites formed deviating size of spheroids. showed spheroid-chain neurite (curved arrow). The diameter of curl-up neurites also presented deviating size. The thicker of the neurites came with larger spheroids and vice versa. It seemed that thinner fibers possessed numerous small size spheroids (Figure 8 B).

**Figure 8.**
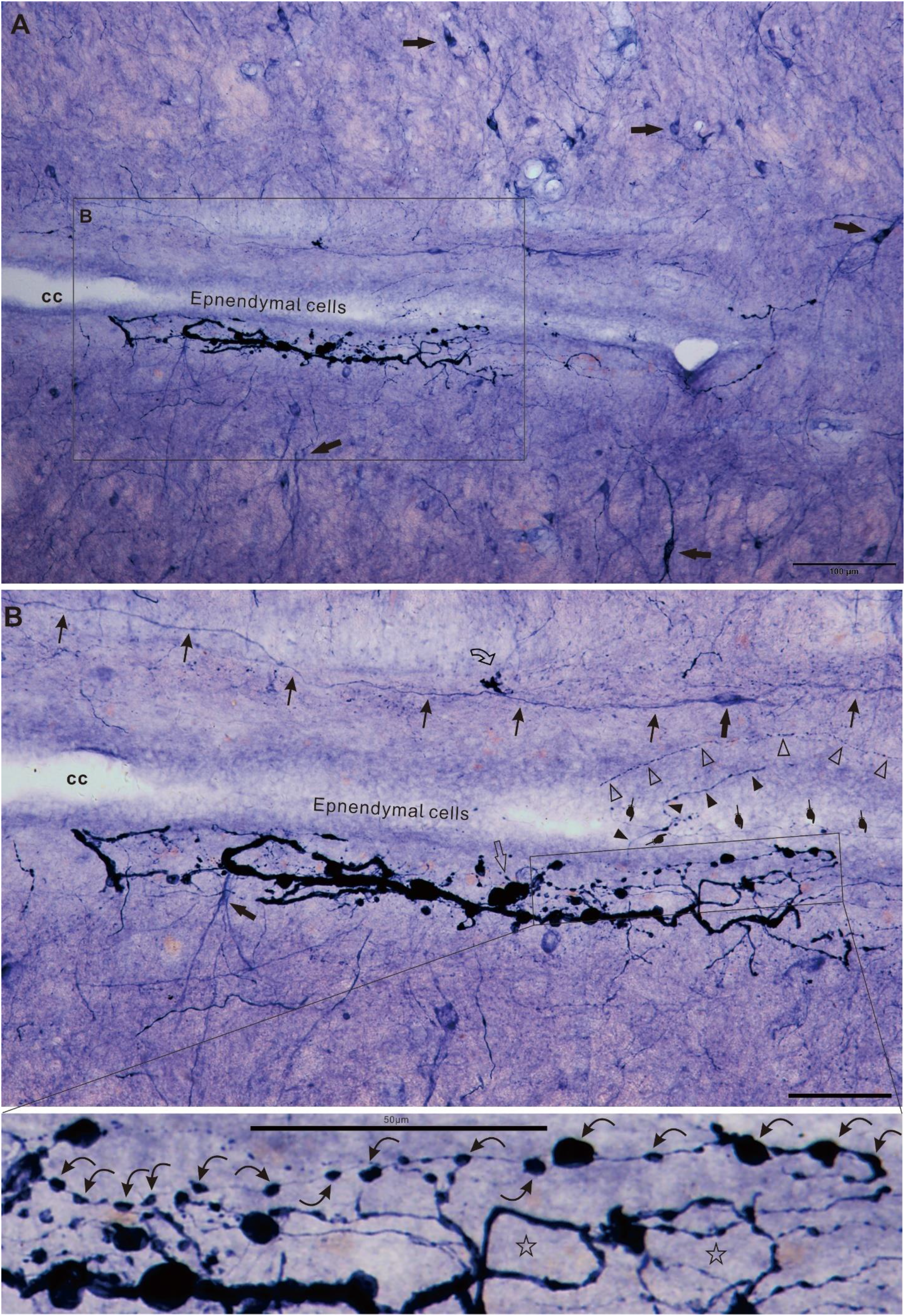
The horizontal section through the central canal showed various diverse neurites(A). Arrow indicated neuron. B magnified from A showed curl-up neurites. Thin arrow: regular fiber. At least three fibers around ependymal cells. Open arrowhead: dash line like fiber. Arrowhead: varicose fiber. Finger point: thin fiber. The bottom image magnified from B showed spheroid-chain neurite (curved arrow). Asterisk indicated circle (coil-like) neurite. Bar in A =100μm. Bar in B and magnified image=50μm.

N-d neurons distribute nearly in all pelvic organ. N-d positive autonomic sensory fibers from dorsal root ganglia (DRG) send central projection into the spinal cord distributed as we demonstrated above. Neurodegenerative pathology occur as vacuolar degeneration in DRG[32]. N-d positive neurons were small size oval neurons (Figure 9). Figure 9 showed vacuolar degeneration in the DRG and thick axons in the rootlet in sacral spinal cord of aged dog. Vacuolar neuron and neuronal shrinkage as well as N-d positive relocation were detected in DRG. N-d fibers were detected in the rootlet and massive N-d fibers in the dorsal white matter. It demonstrated that substantial neurodegeneration essentially occurred in the spinal cord.

**Figure 9.**
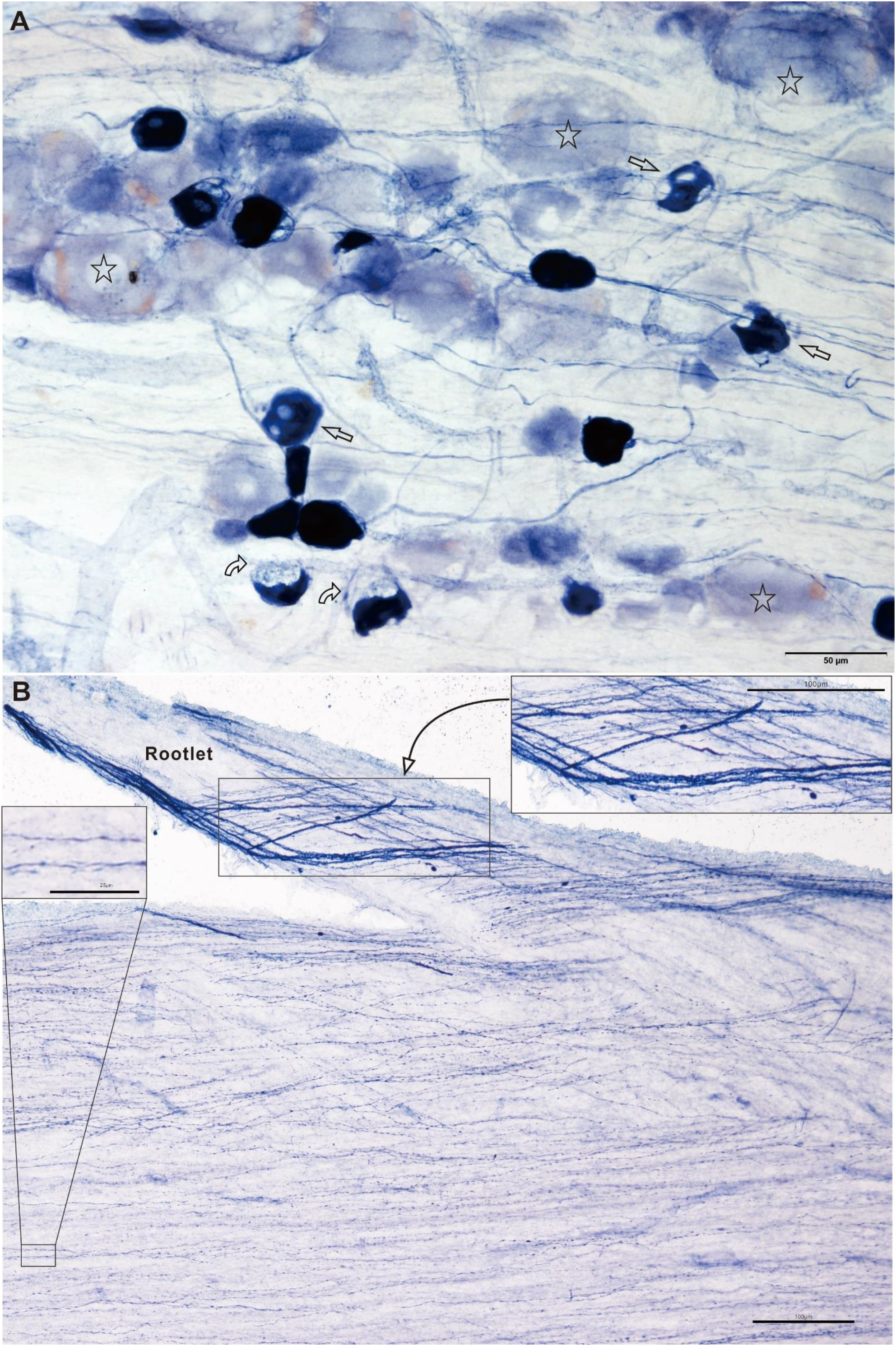
Vacuolar degeneration revealed in the dorsal root ganglia (DRG) and thick axons in the rootlet in sacral spinal cord of aged dog. A showed vacuolar neuron (open arrow) in DRG. Curved open arrow indicated. Curved open arrow indicated N-d positive relocation and neuronal shrinkage. B showed N-d fibers in the rootlet and massive N-d fibers in the dorsal white matter. Bar in A =50μm, B and inset 2=100μm, inset 1 = 25μm.

No typical megaloneurites were detected in the lumber segments. Irregular dystrophic neurite, mini spheroid and ANB were revealed in the dorsal horn (Figure 10). Eccentric dystrophic neurites revealed spheroid and ANB, showed as mini spheroid and large size spheroids, although no typical megaloneurites detected in dorsal horn between the lateral funiculus (LF) and dorsal column.

**Figure 10.**
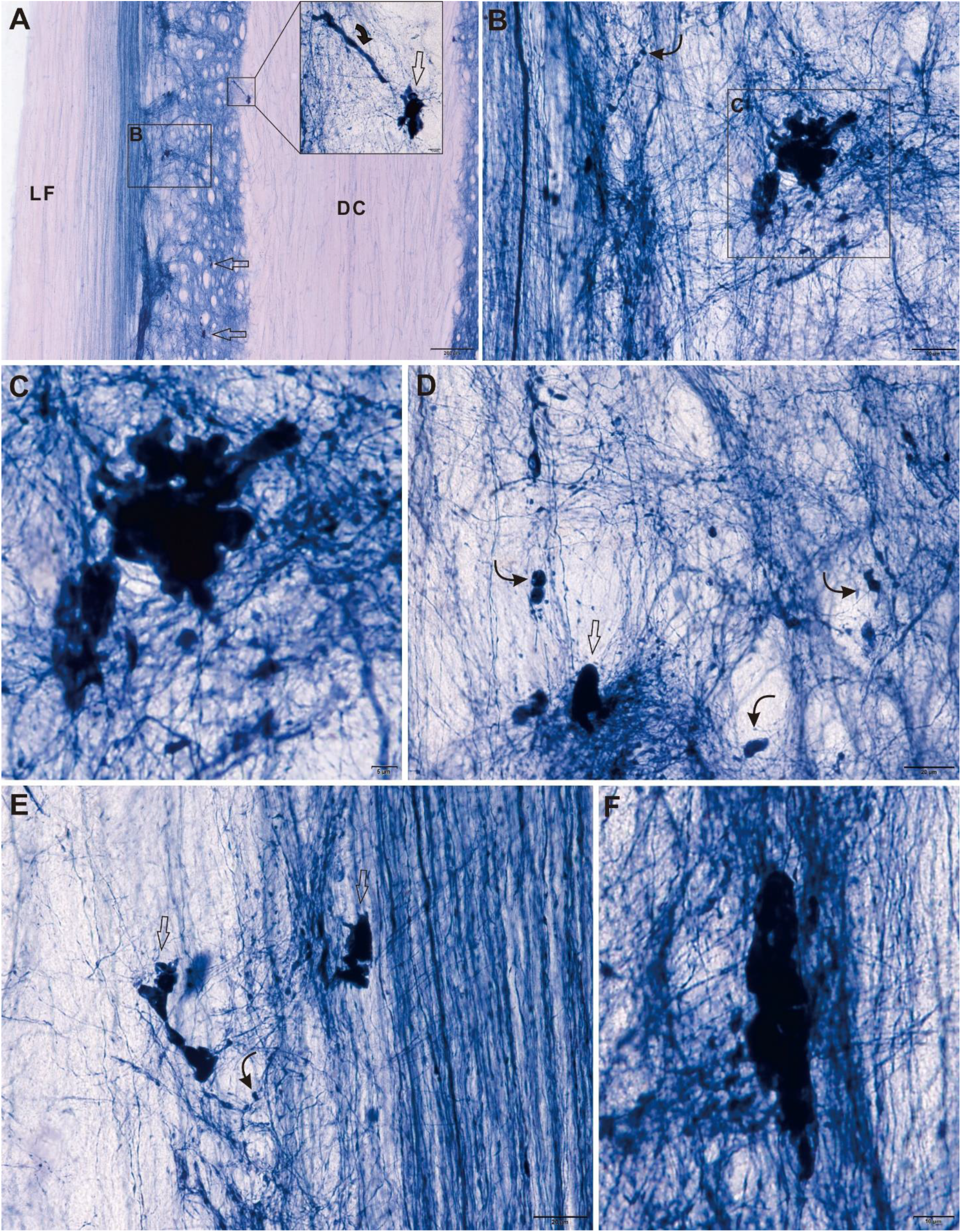
Horizontal section for L5 through the dorsal horn. A-C reviewed from the same microscope field. No typical megaloneurites detected in dorsal horn between the lateral funiculus (LF) and dorsal column. Eccentric dystrophic neurites revealed in A-C. Curved arrow indicated spheroid and ANB. D-F similar location as A-C showed mini spheroid (curved thin arrow) and large size spheroids (open arrow). Bar in A=200μm, B, D and E=20μm, C= 5μm, F= 10μm.

Parasympathetic nucleus locates in the intermediate regional and or IML in the sacral spinal cord. The curl-up and coiled neurites occurred in the regions in the old dog. The corresponding location of parasympathetic nucleus for sympathetic nucleus locates in the upper lumber and thoracic spinal cord. Figure 11-1 showed that the ANB and vacuolar neurite occurred in the horizontal section at the lumbothoracic spinal cord. With study of adrenal innervation, Cummings subdivide autonomic neurons into four groups, central autonomic neurons, base of ependymal cells at dorsal border of central canal, intercalated cells, nucleus intermediolateralis for the thoracolumbar preganglionic neurons and adrenal innervation in the dog[33]. In Figure 11, we examined the spatial localization of IML in typical horizontal section through the central canal indicated by ependymal cell (EC). The preganglionic neurons formed the typical IML in the thoracolumbar spinal cord. The ANBs and a thick proximal dendritic neuron revealed in Figure 11-1. Numerous varicose fibers and punctus also distributed around the dorsal area of central canal. Similar data continued to show the base of ependymal cell dorsal of the central canal (Figure 11-2).

**Figure 11-1.**
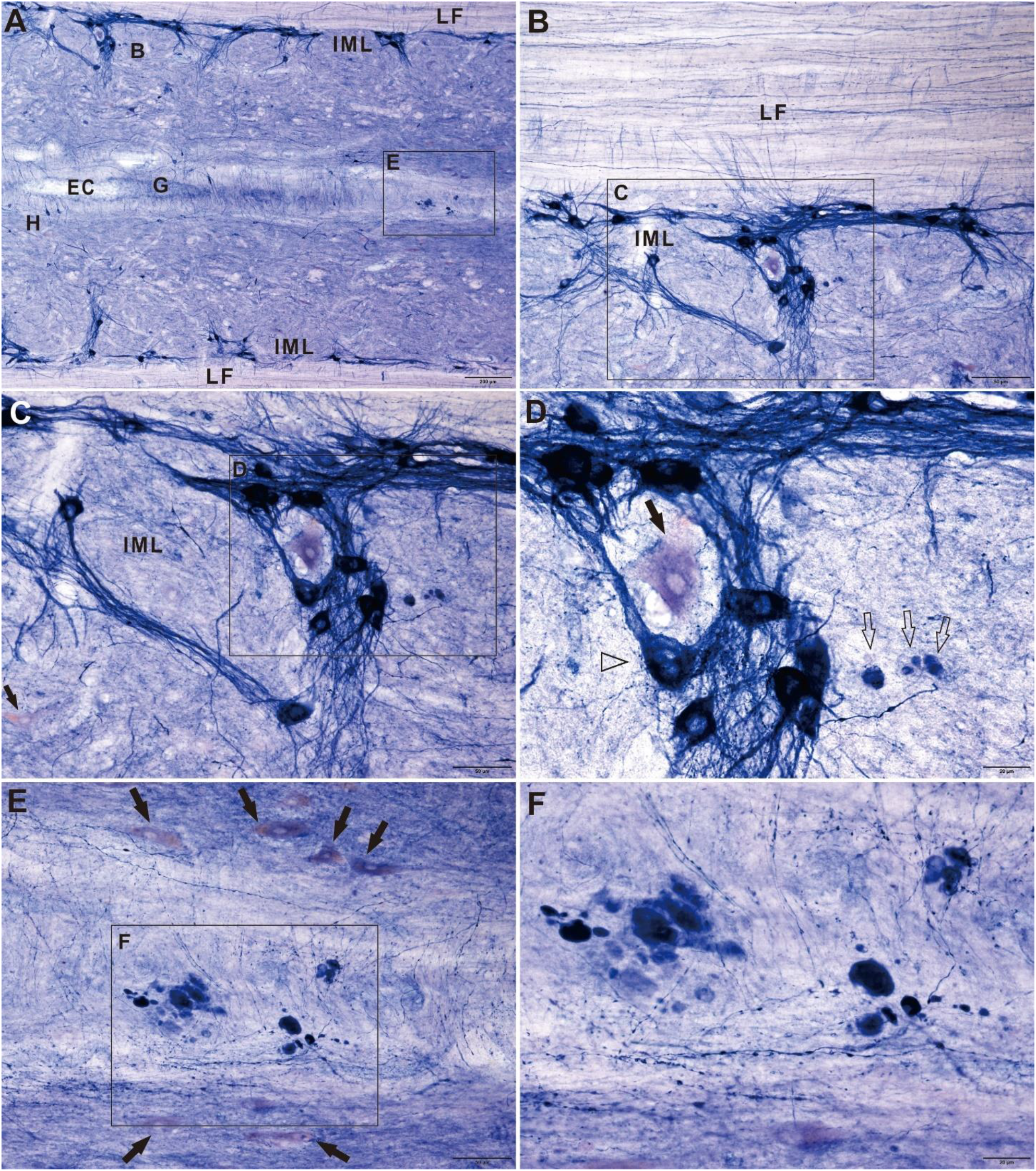
The ANB and vacuolar neurite occurred in the horizontal section at the lumbothoracic spinal cord. A: The spatial localization of IML in typical horizontal section through the central canal indicated by ependymal cell (EC). B, E, G and H labeled for following magnified images. LF indicated the lateral funiculus. B, C and D demonstrated the ganglionic cells and fibers in IML. Arrow indicated an unstained neuron in C and a large lightly stained neuron surround by the ganglionic neurons in D. The ANBs (open arrow) and a thick proximal dendritic neuron (open arrowhead) revealed in D. E and F showed cluster of ANB as well as neurons (arrow) in E. Varicose fibers and punctus distributed around the dorsal area of central canal. Bar in A =200μm, B=100μm, C and E=50μm, D and F =20μm.

**Figure 11-2.**
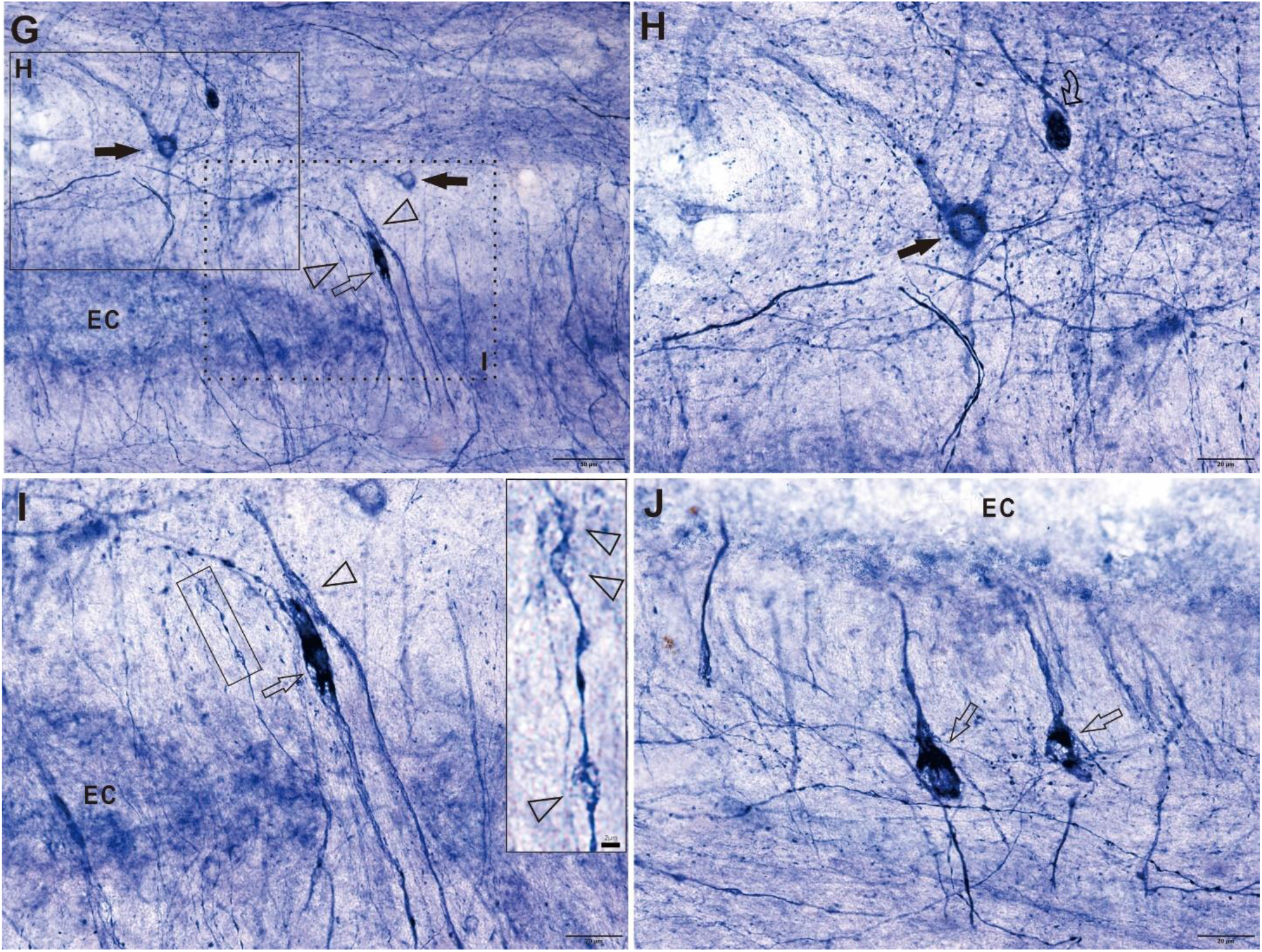
Continued with the Figure 11-1 to show the base of ependymal cell (EC) dorsal of the central canal. All images magnified from Figure 11-1A. Similar to Figure 11-1E and F, numerous of varicose fibers and puncta distributed in G-J. H showed a neuron (arrow) and ANB. I magnified from dash line rectangle of G showed vacuolar swelling neurite (open arrow) and similar dilation fiber (open arrowhead). Inset showed a bulb-like dilated fiber (open arrowhead). J showed two vacuolar neurons. Bar in G =50μm, H-J 20μm and inset = 2μm.

## DISCUSSION

In the present study, megaloneurites was consistently confirmed with our previous investigation [24]. With N-d histology, the horizontal section of the longitudinal fibers and tracks provided new dimensional view to traditional architecture of the spinal cord. The main result of the present investigation indicated that N-d positive mini spheroid, regular spheroid occurred in the marginal laminae and spacing curl-up and coiled neurites specifically located intermediate lateral region in the sacral spinal cord of aged dog. The coiled neurites showed spacing arrangement which similar to chain pattern of distribution of the parasympathetic preganglionic neuron. The individual coiled neurites also sent fiber bundle to the DCN (Figure 7-1B). The longitudual neurites varied from thin and thick fibers as well as megaloneurites. The pathway of the all neurites revealed fasciculus proprius and or cross segment funiculus. Acording to Figure 2 to 5 and Figure 9, the megaloneurites occurred in the LT, dorsal lateral funiculus, MCP and LCP. The Figure 5-2C and Supplementory Figure 1 showed that spheroids (open arrow) located in focal regions of swelling neurite. It could be speculated that the N-d positivity acumlated in the neurite and the acumlation cause neurites dilation and blokade of normal neuritic transportation. Figure 5-2F showed inset of compound spheroid with relative shallow staining halo surround high intensity core. It could be speculated that the N-d leaked from the swelling neurites. The vacuolar neurons and neurites revealed in the thoracic, lumber and sacral spinal cord. This was similar other neuropathology of neurodegenerative diseases [34–39]. It could be speculated that the aging-related N-d alteration was neuropathological degeneration. Summarily, we supposed that the spacing curl-up neurites and coiled neurites may be neurodegeneration of autonomic input of N-d positivity in the sacral spinal cord which caused the autonomic circuit dysfunction of pelvic organs.

Combined with our previous report of disc herniation of aged dog[25] and previous report of megaloneurite[24], aging deterioration of N-d positivity in the sacral spinal cord showed selective destruction of N-d neuronal elements, the pathway of N-d input to the sacral spinal cord and spacing pattern of projecting terminals. The spacing mini spheroids in the laminae I of the dorsal horn was similar to the neuropil spheroids in the neuropathological changes of the frontotemporal dementia [40]. For the pattern of spacing curl-up and coiled neurites, we resumed neurodegeneration of spacing neuronal wire and circuit for autonomic modulations, impaired neuronal circuit of dorsal part of the spinal cord.

For small-fiber sensory neuropathies, Holland describes that the swelling or spheroidal alteration of nerve terminals also occur in the somatic sensory dysfunction, besides abnormally segmented terminals[41]. In our research results, N-d spheroids, especially small-size spheroids distributed in the dorsal horn. But, N-d neurons and neuropils also distributed in the ventral horn. In some neurodegenerative diseases, neurodegenerative spheroids distribute in the anterior horn[40].

The aging causes the degeneration of spinal cord. “the normal fibers appeared as thin, unbroken profiles”, while the criteria of degenerating fibers are the beaded and short, rod-like and fiber breakdown appearance[42]. The N-d positive LCP normally terminates IML region. IML can be sub-grouped. Retrograde neuronal tracing study reveals the nucleus intermediolateralis inferior in the sacral autonomic neurons in the cat. The neurons of the intermediolateral group are also arranged in horizontal fashion [43]. Petras and Cummings demonstrate that the bladder and urethra are innervated by several nuclear groups. The bladder receives fibers from two lumbar (the nucleus intermediolateralis thoracolumbalis pars principalis, ILp; nucleus intercalatus spinalis, IC) and from 3 sacral (parasympathetic nucleus intermediolateralis sacralis pars principalis, ILSp; nucleus intcrmediolateralis sacralis pars funicularis, ILSf and nucleus intermediolateralis sacralis pars ala ventralis, ILSav) nuclei, while the urethra receives contributions from 4 sacrococcygeal nuclei (ILSp, ILSf, ILSav and nucleus intercalatus disseminate, ICd) neonatal dogs [10]. Both the spacing curl-up neurites and coiled neurites located the same regions of IML and intermediate zone.

The vacuolated neurodegeneration in the present was consistent with our previous case report of disc herniation[25]. Megaloneurite, ANB or N-d spheroid are relatively specific neurodegeneration in sacral spinal cord of aged dog. The vacuolated neurodegeneration is commonly detected in other neurodegenerative diseases. For example, ballooned neurons with distended, vacuolated perikarya and or proximal neuronal processes are considered as histological features of several neurodegenerative diseases of the central nervous system [34]. Neuritic bulbs or fragmentation is also commonly feature of neurodegeneration[35] [36]. Distal dendritic expansion is pathological appearance of neurodegeneration of Tauopathy for extracellular deposit [37]. Autonomic dysfunctions of orthostatic hypotension, urinary incontinence and constipation are common symptoms in neurodegenerative demented patients [38]. Neurodegeneration of the sacral spinal cord may be relevant the onset of multiple system atrophy[39]. Onset of urinary retention in multiple system atrophy can begin in the sacral spinal cord [39]. Ballooned neurons can occur in the IML of the sacral spinal cord[44]. We noted that early cauda equina syndrome of a model of somatovisceral pain in dogs may not involve in vacuolar perikarya, axonal bulbs or fragmentation with N-d histology in spinal cord structures [45]. Aging-related or chronic alteration may be important risk causative factors for normal aging deterioration or neurodegenerative diseases. An acute model may not cause dramatically on the aging change spinal cord.

Central autonomic neurons, base of ependymal cells at dorsal border of central canal, intercalated cells, nucleus intermediolateralis. Thoracolumbar preganglionic neurons and adrenal innervation in the dog [33]. Intermediolateral nucleus located in the lateral horn of thoracic [46] and lumbosacral spinal cord [47]. The innervation of pelvic floor muscles and bladder neck in the dog reveals that most of spinal cord neurons distribute in the ventral horn, however, some neurons scatter at the dorsal lateral to the intermediate zone close to IML in the level of the central canal in the sacral spinal cord [9]. We noted that Lanerolle et al use “cell nests” to further define the location of IML or use “terminal-like processes”, “terminal-like processes”, “terminal field” and “terminal-like structures” to describe distribution of substance P-like and methionine-enkephalin-like immunoreactivity [48]. We may use “dystrophic neuritic nest” or “dystrophic neuritic plexus” to describe the format or profile of the curl-up neurites and coiled neurites. For control of the micturition process, the several routes relate a closed loop system[49, 50]. N-d neurons involve the autonomic input fibers and preganglionic output.

Aging-related urogenital dysfunctions may be corelated with the circuit wiring disorders of neurite terminal arborization. The functional organization of N-d specific intrinsic circuitry was specifically disorganized for the spheroids, spacing coiled neurites and previously reported megaloneurite as well as homogeneous formazan globule. We supposed that those results were a new dimension of aging neurodegenerative alteration in the sacral spinal cord of aged dog. Incontinence, constipation, erectile dysfunction[51] and pelvic organ prolapse are aging-related phenotypic manifestations[52, 53]. Neuro-autonomic regulation is considered with pathophysiology of urinary incontinence, fecal incontinence and pelvic organ prolapse[54–56]. Importantly, N-d-ergic innervation is considered relevant to pelvic function and disorders[57–62]. When the curl-up neurites and coiled neurites became aged aberrant, the function of urogenital organs faltered. The N-d innervation in the sacral spinal cord became a vulnerable part with aging deterioration, to partake the incontinence, constipation, erectile dysfunction and pelvic organ collapse. Bulterijs et al demonstrate biological aging to classify as a disease[63]. We further hypothesis that the abnormality of the N-d positivity in the sacral spinal cord and dysfunction of urogenital organs agree with “biological aging as a disease”. We concluded that N-d staining could visualized specific neuropathology. Some morphological criteria of megaloneurite, ANB, spheroid, curl-up neurite and coil neurite are conceived as sacral spinal cord specification. Vacuolized neurodegeneration is consistent with neuropathology of common neurodegenerative diseases.

## ACKNOWLEDGMENTS

This work was supported by grants from National Natural Science Foundation of China (81471286) and Research Start-Up Grant for New Science Faculty of Jinzhou Medical University (173514017).

---

**Supplementary Figure 1.**
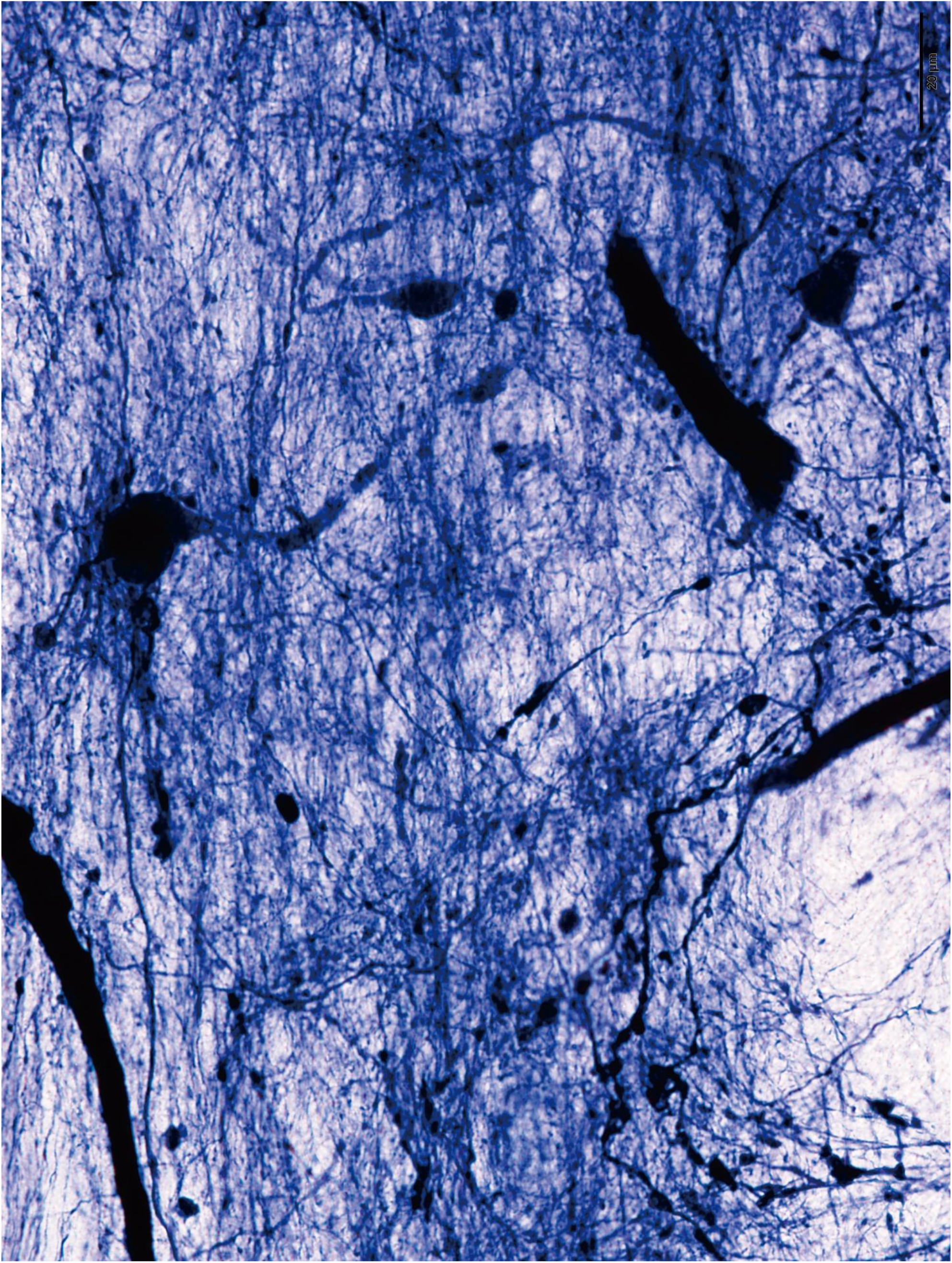
A large format image of the same photo image of Figure 5C. Bar=20μm

**Supplementary Figure 2.**
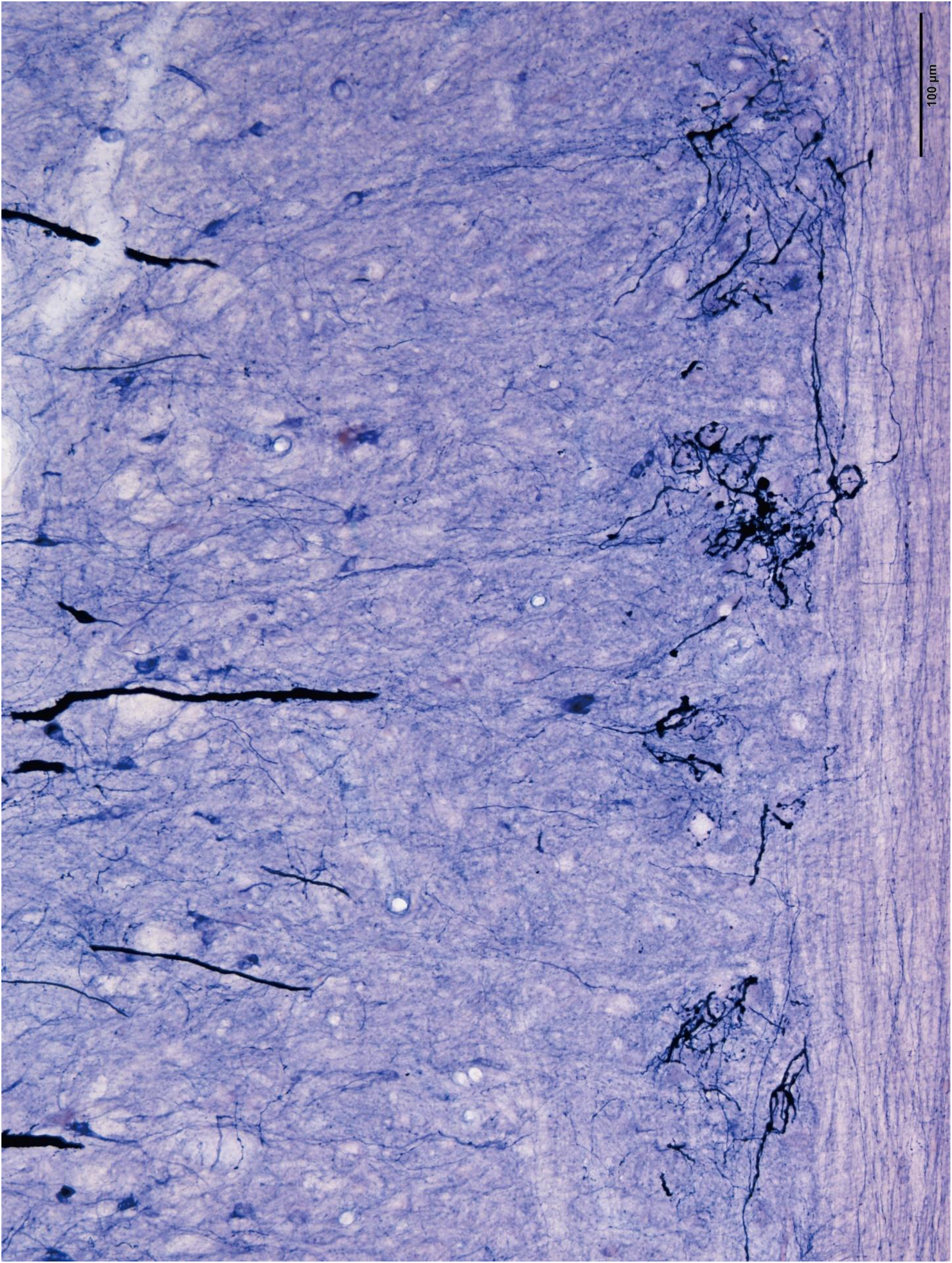
A large format image of the same photo image of Figure 7-1B. Bar=100μm

**Supplementary Figure 3.**
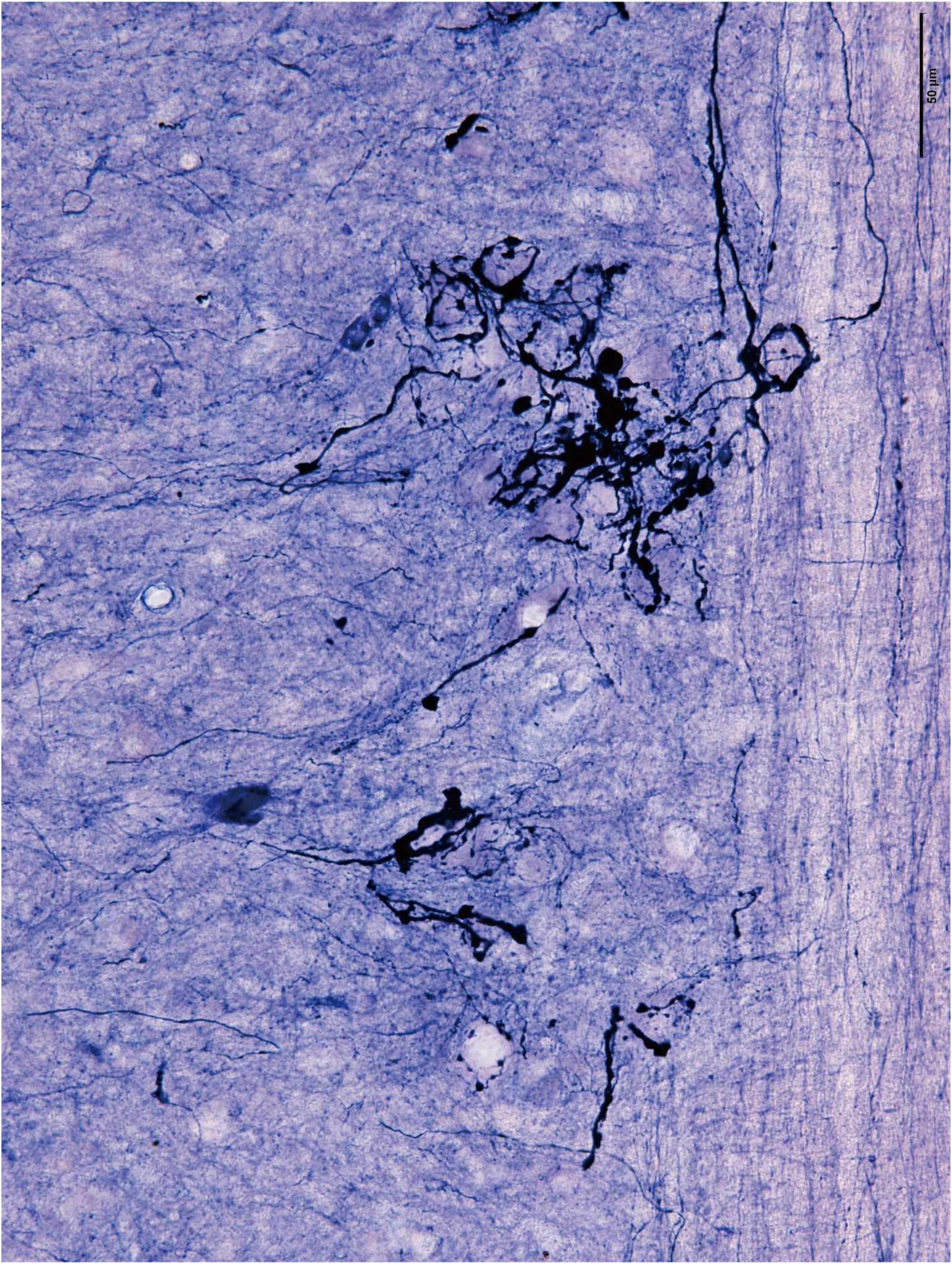
A large format image of the same photo image of Figure 7-1C. Bar=50μm

